# Investigation of an *LPA* KIV-2 nonsense mutation in 11,000 individuals: the importance of linkage disequilibrium structure in *LPA* genetics

**DOI:** 10.1101/848945

**Authors:** Silvia Di Maio, Gertraud Streiter, Rebecca Grüneis, Claudia Lamina, Manuel Maglione, Dietmar Öfner, Barbara Thorand, Annette Peters, Kai-Uwe Eckardt, Anna Köttgen, Florian Kronenberg, Stefan Coassin

**Author notes:** **Address for correspondence**, Stefan Coassin, PhD, Institute of Genetic Epidemiology, Department of Genetics and Pharmacology, Medical University of Innsbruck, Schöpfstrasse 41, A-6020 Innsbruck, AUSTRIA.

## Abstract

**Objective:** Elevated Lp(a) plasma concentrations are determined mainly genetically by the *LPA* gene locus, but up to 70% of the coding sequence is located in the so-called “kringle IV type 2” (KIV-2) copy number variation. This region is not resolved by common genotyping technologies and large epidemiological studies on this region are therefore missing. The Arg21Ter variant (R21X, variant frequency ≈2%) is a functional variant in this region, but it has never been analyzed in large cohorts and is it unknown whether it is captured by genome-wide association studies.

**Approach and Results:** We developed a highly sensitive allele-specific qPCR assay and genotyped R21X in 10,910 individuals from three populations (GCKD, KORA F3, KORA F4). R21X carriers showed significantly lower mean Lp(a) concentrations (−11.7 mg/dL [−15.5;−7.82], p=3.39e-32). Of particular note, virtually all R21X carriers also carried the splice mutation rs41272114 (D’=0.957, R^2^=0.275), as confirmed by pulsed-field gel electrophoresis and long-range haplotyping. This proposes that the R21X mutation arose on the background of the rs41272114 splice variant.

**Conclusions:** We performed the largest epidemiological study on an *LPA* KIV-2 variant so far. Interestingly, R21X is located on the same haplotype as the splice mutation rs41272114, creating “double-null” *LPA* alleles that are inactivated by two independent mutations. The effect of the R21X nonsense mutation can thus not be discerned from the effect of rs41272114 splice site mutation. This emphasizes the importance of assessing the complex LD structure within *LPA* even for functional variants.

## Introduction

The lipoprotein(a) [Lp(a)] plasma concentration is one of the most prominent genetically determined factors for cardiovascular diseases^1–7^. Elevated Lp(a) concentrations concern 14-25% of the general population^2^ and increase the cardiovascular disease (CVD) risk up to more than three-fold for very high concentrations above the 99^th^ percentile^8^. More than 90% of Lp(a) variance is genetically determined by the *LPA* gene locus^9^, which encodes apolipoprotein(a) [apo(a)], the distinctive structural protein of the Lp(a) particle. The apo(a) protein consists of 10 types of so-called kringle-IV domains (KIV-1 to −10), one kringle V domain and a protease domain. The KIV-2 domain is encoded by a hypervariable copy number variation (CNV), which is present in up to ≈40 copies per gene allele, resulting in a pronounced size polymorphism of the apo(a) protein^10^ and theoretically ≈1600 possible genotypes in the population. The KIV-2-CNV explains 30-70% of the Lp(a) _1_ concentrations.

The apo(a) isoform size is inversely correlated with Lp(a) concentrations^1^. Low molecular weight (LMW) apo(a) isoforms (≤22 KIV domains) are associated with 5-10 fold higher median Lp(a) concentrations than high molecular weight (HMW) isoforms (>22 KIV domains)^1^ but the Lp(a) levels of unrelated individuals carrying a same-sized isoform combination can still vary by up to 200-fold^11,12^. In contrast, same-sized alleles within families (i.e. alleles that are identical-by-descent) are associated with a much smaller variation in Lp(a) (typically <2.5-fold^11^). This indicates that genetic variants exist that regulate the Lp(a) concentrations in addition to isoform size ^12,13^. Genome-wide association studies (GWAS) have identified dozens of SNPs distributed over a two megabases region^14–16^, but few bona-fide *functional* variants have been identified^13,17–19^.

The KIV-2 region encompasses up to 70% of the *LPA* coding sequence^13^ and is therefore an obvious candidate region to search for functional variants affecting Lp(a) concentrations. Very little is known about the impact of sequence variation in the KIV-2 CNV on Lp(a) concentrations because commonly used sequencing and genotyping technologies are not able to resolve variation within this region. Accordingly, it is also unclear whether KIV-2 variants are captured by known GWAS hits via linkage disequilibrium (LD) or whether they are indeed mostly independent. Some studies suggested a strong LD structure spanning the whole kringle region^20–23^, but few details have been reported.

Recently, an ultra-deep next generation sequencing (NGS) approach with a customized bioinformatic analysis pipeline has allowed cataloging variation within the KIV-2 region^24^. This revealed hundreds of variants and provided several new putative regulators of Lp(a) levels. For example, the G4925A variant^13^ found in 20% of the population is associated with an Lp(a) reduction of up to ≈30 mg/dL and explains a considerable fraction of the individuals presenting low Lp(a) concentrations despite carrying a LMW isoform. This aspect of the relationship between Lp(a) concentrations and apo(a) isoform size is poorly understood so far.

The nonsense mutation KIV-2 R21X (g. 61 C>T in ^19^, 640C>T in ^24^) is another likely causal single nucleotide polymorphism (SNP) in the KIV-2 region. It leads to a truncated protein that is rapidly degraded^19^. Parson et al.^19^ identified it by a laborious cloning approach and reported a minor allele frequency [MAF] of 1.67% in 405 individuals^19^. However, because standard genotyping technologies like TaqMan genotyping assays and SNP microarrays are not specific and sensitive enough to detect a variant that is present in only one (or a few) of up to 80 nearly identical repeats (i.e. 1.2% mutation level), the R21X variant was not further explored in large epidemiological studies. Accordingly, R21X has never been put into the context of the findings from GWAS on Lp(a)^14–16^ and it is unknown whether any of the *LPA* SNPs detected in GWAS studies is in LD with R21X.

We developed an allele-specific TaqMan PCR assay (ast-PCR) targeting the R21X variant, as well as the previously described KIV-2 variant G4925A^13^, and assessed the effect of R21X on Lp(a) concentrations in nearly 11,000 individuals. We then used genome-wide SNP data from the German Chronic Kidney Disease (GCKD) study to assess the LD of R21X with SNPs outside the KIV-2 region to link it to available GWAS datasets. Finally, given the fact that the effect of a functional *LPA* SNP depends strongly on which gene allele it is located, we determined the allelic location of R21X by pulsed-field electrophoresis (PFGE) and show the existence of moderately frequent “double-null” *LPA* alleles that are inactivated by two independent loss-of-function mutations.

## Materials and Methods

### Populations

Our study involved 10,910 individuals from three studies, namely GCKD^25^ (German Chronic Kidney Disease), KORA (Cooperative Health Research in the region of Augsburg) F3^26^ and KORA F4^26^. Informed consent was obtained from each participant and the studies were approved by the respective Institutional Review Boards. Details on the studies are given in Table 1 and in the Supplemental Methods. In brief, the KORA F3 and F4 studies are follow-up examinations of two nonoverlapping surveys drawn from the general population living in the region of Augsburg, Southern Germany (KORA S3 and S4). GCKD is an ongoing prospective observational study of 5,217 Caucasian patients with moderately severe chronic kidney disease at enrollment recruited at nine institutions in Germany.

**Table 1.** Descriptive statistics. Continuous variables are provided as mean ± SD and [25%, 50%, 75%] percentiles. LMW: low molecular weight apo(a) isoforms. HMW: high molecular weight apo(a) isoforms. F: females. M: males. eGFR: estimated glomerular filtration rate. Lp(a): lipoprotein(a).

|  | GCKD | KORA F3 | KORA F4 |
| --- | --- | --- | --- |
| n | 4,771 | 3,099 | 3,040 |
| Sex (F) | 1,896 (39.7%) | 1,602 (51.7%) | 1,576 (51.8%) |
| Age (years) | 60.1±11.9<br>[53,63,70] | 57.4±12.9<br>[46,57,67] | 56.1±13.3<br>[44,56,67] |
| Age (F) (years) | 59.0±12.7<br>[51,63,69] | 57.1±12.7<br>[47,57,67] | 55.6±13.1<br>[44,56,67] |
| Age (M) (years) | 60.9±11.3<br>[55,64,70] | 57.6±13.1<br>[46,58,68] | 56.6±13.4<br>[45,57,68] |
| Lp(a) (mg/dL) | 24.7±30.6<br>[5.0,11.6,33.7] | 22.21±26.2<br>[5.0,11.1,29.1] | 21.7±24.6<br>[5.2,11.7,30.23] |
| Lp(a) in LMW (mg/dL) | 56.5±40.4<br>[25.7,53.2,77.9] | 50.6±34.2<br>[18.3,49.4,72.6] | 47.8±31.2<br>[23.7,46.8,66.5] |
| Lp(a) in HMW (mg/dL) | 14.3±16.3<br>[3.8,8.59,17.8] | 13.6±14.8<br>[4.02,8.75,16.7] | 13.4±14.1<br>[4.11,8.95,17.3] |
| eGFR (mL/min/1.73 m<sup>2</sup>) * | 49.4±18.2<br>[37,46,57] | 86.7±17.5<br>[76.14,89.11,99.2] | 87.84±16.9<br>[77.4,89.3,100.1] |
| Cholesterol (mg/dL) | 211.1±52.7<br>[176.5,207.1,239.1] | 218.1±39.9<br>[191,216,242] | 216.0±39.6<br>[188,214,240] |
| HDL-C (mg/dL) | 51.8±18.0<br>[39.21,48.24,61.15] | 58.77±17.2<br>[46,56,69] | 55.9±14.4<br>[45,54,65] |
| LDL-C (mg/dL) | 118.4±43.6<br>[89.2,113.8,142.7] | 127.88±32.6<br>[105,126,148] | 136.07±34.9<br>[112,134,158] |
| Triglycerides (mg/dL) | 198.2±125.0<br>[117.6,168.0,238.7] | 164.85±125.1<br>[88,135,200] | 124.71±88.9<br>[71.5,104,150] |
| Type 2 diabetes mellitus | 1,713 (35.9%) | 248 (8.1%) | 241 (7.9%) |

\* Estimated using the CKD-Epi equation<sup>39</sup>

DNA specimens from common study populations are not suitable for pulsed-field gel electrophoresis (PFGE), because it requires megabase-sized DNA. Since the buffy coat samples that are required for such an agarose-plug DNA preparation are not commonly available for population studies, we used for the PFGE experiments samples from the CAVASIC study^27,28^, from an ongoing collection of liver tissue specimens for Lp(a) research (IRB Medical University of Innsbruck, AN2015-0056) and from anonymous blood samples obtained from the blood bank of the University Hospital of Innsbruck, Austria.

### Ast-PCR for R21X typing

We designed an allele-specific triplex TaqMan PCR assay (ast-PCR) amplifying the mutant bases of R21X^19^ and the G4925A^13^, as well as a positive amplification control (design illustrated in Supplemental Figure I). The R21X and the G4925A variants were introduced into pSPL3 plasmids containing one KIV-2 repeat^13^ using the Agilent Technologies QuickChange II Site-Directed Mutagenesis Kit with minor modifications, Supplemental Methods). To provide additional thermodynamic disadvantage to unspecific pairings ^29^, various base mismatches were introduced in the allele-specific primers on positions −2 or −3 (from the 3’ end) and the performance of different primer designs was tested on plasmid mixes mimicking mutation levels from 100% to 0% mutant fraction (Supplemental Figure II). The most specific primers were taken forward (Supplemental Figure II). Fluorescently labelled, locus-specific TaqMan probes were added to allow amplification detection in a high throughput setting. An amplicon in *PNPLA3* served as positive amplification control to detect false-negative reactions due to PCR failure. Technical details are provided in the Supplemental Methods and in Supplemental Tables I and II. The assay was run on a 384-well ThermoFisher QuantStudio 6 qPCR system. The R21X assay was validated both against ultra-deep NGS data from Coassin and Schönherr et al, 2019^24^ and against a commercial cast-PCR assay (ThermoFisher; as used in ^13^) with a sensitivity of 0.2% mutant fraction (determined according to manufacturer instructions on a NGS-validated sample). For validation, our assay was run on 376 samples from KORA F4, identifying 14 R21X-carriers, which were all confirmed also by the commercial castPCR assay. The reproducibility was tested on 477 samples run in duplicates. Additionally, each 384 well qPCR plate (n=34) contained a positive control sample. A slightly modified ast-PCR protocol was used to genotype the gene alleles separated by PFGE (Supplemental Methods).

### Ast-PCR data analysis

Figure 1 exemplifies the assay data analysis rationale. A possible unspecific amplification signal of the WT variant alleles will occur later in the qPCR amplification than the true specific amplification of the mutant variant allele [C_T(carriers)_<C_T(non-carriers)_]. This creates two C_T_ distributions whose widths are defined by stochastic fluctuation in the amplification of the target (e.g. due to slightly varying input amount) and, for the mutant, the fraction of KIV-2 repeats affected (Figure 1A). DNA input was 20 ng in all samples. Previous data^13,24^ indicates that no more than one to three repeats are affected by R21X, which translates to maximum ≈1.6 cycles differences due to the mutation level.

**Figure 1.**
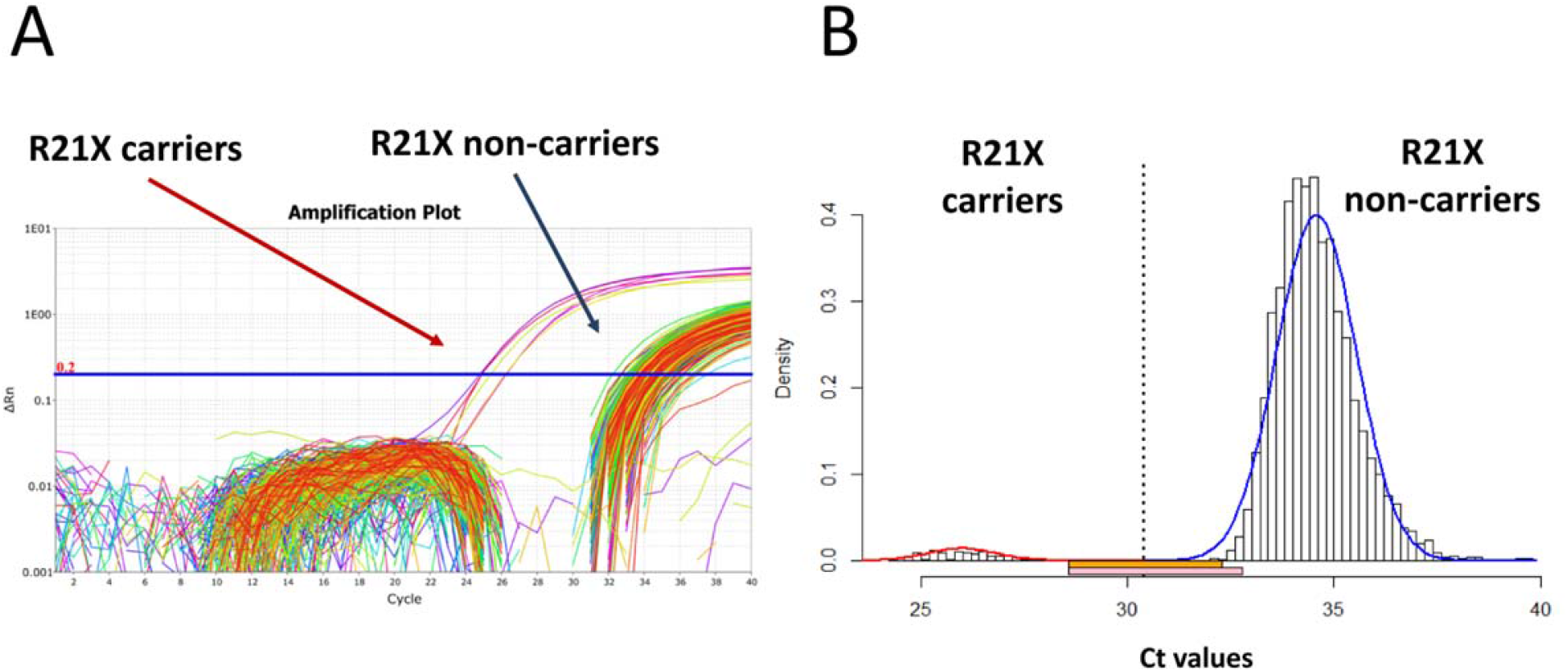
Analysis strategy. **A.** Exemplary ast-PCR amplification plot. **B.** Discrimination of the two C_T_ distributions using a systematic clustering approach. The orange and pink horizontal bars (below the x-axis) identify the samples, which cannot be uniquely assigned to one of the two distributions and are therefore excluded from the analysis. Orange bar: upper 1% of the carrier distribution and lower 1% of the non-carrier distribution. Red bar: upper 2.5% of the carrier distribution and lower 2.5% of the noncarrier distribution (more conservative; used in this analysis). Plot generated using the provided R script.

To avoid human bias and have a systematic approach for sample assignment beyond pure visual clustering of the amplification curves, the optimal discrimination threshold between the CT distributions of carriers and non-carriers was estimated using a bagged clustering algorithm^30^ implemented in the R function ‘classIntervals’ (package ‘classInt’) and two normal distributions were fitted to the two CT distributions using the R package ‘VGAM’^31^ (Figure 1B). Details are provided in the Supplemental Methods. Samples that could not be assigned unambiguously to one of the CT distributions (i.e. which cannot be unambiguously identified as carriers or non-carriers) were excluded. The exclusion rate was 0.7% in GCKD (35/4974), 1.6% in KORA F3 (52/3157) and 0.5% in KORA F4 (15/3063). The custom R function used for analysis is available at https://github.com/scogi/r21x_analysis.

### Lp(a) phenotyping

The Lp(a) concentrations and apo(a) isoforms were determined by ELISA and Western blotting, respectively, as described earlier^32,33^. All analyses were performed in the same laboratory at the Institute of Genetic Epidemiology, Medical University of Innsbruck, Austria and evaluated by the same experienced researcher.

### Identification of proxy SNPs

We used the genome-wide SNP data available for the GCKD study to search for a proxy SNP for R21X that would allow linking R21X to existing results from GWAS. Following the rationale that a SNP in LD with R21X will present a similar effect on Lp(a) and that the observed effect of R21X on Lp(a) should have been easily detected by our recent GWAS on Lp(a) (n=13,781)^14^, we created a contingency table of each of the 66 top hits of the isoform-adjusted model of our recent GWAS metaanalysis on the Lp(a) concentrations with the R21X^14^ and analyzed it using the Fisher’s exact test. We selected the two SNPs with the most significant p-values (rs2489940, rs41272114) and calculated the LD using CubeX^34^.

### Pulsed-field gel electrophoresis

We performed *LPA* PFGE-based genotyping^13,35–37^ to assess on which gene allele R21X is located and to confirm the co-localization of R21X and rs41272114 experimentally. Two different restriction enzymes were used for *LPA* PFGE. KpnI excises a region from KIV-1 to KIV-5^35–37^ and allows precise allele sizing. Conversely, Kpn2I digestion excises a much larger fragment that spans from *LPA* to *MAP3K4*^38^ (Supplemental Figure III) and allows long-range haplotyping of variants by performing the genotyping directly on the previously separated gene alleles.

*The LPA* gene alleles of eight R21X carriers and three R21X-negative samples were separated by PFGE and detected by Southern blotting using a probe against KIV-2 (detailed in ^36^ and ^38^). In brief, DNA agarose plugs have been prepared as previously described^38^ and digested for 4 hours at 37°C (KpnI) or 55°C (Kpn2I). Half plug for each sample was applied on the agarose gel twice, separated on a Bio-Rad CHEF Mapper system (for technical details see Supplemental Table III) and the separated alleles were isolated from the gel as described previously^13,36^. DNA was extracted from the gel slices using the peqGOLD Gel Extraction Kit (VWR). Genotyping was done using a modified ast-PCR protocol for R21X (Supplemental methods) and Sanger sequencing for rs41272114 (Supplemental Table IV).

### Statistical methods

Differences in medians were assessed by Wilcoxon tests. The association between the *LPA* KIV-2 variant R21X and the Lp(a) levels was assessed by linear regression analysis in each population, adjusted for age and sex. GCKD analysis was also repeated adjusting additionally for the estimated glomerular filtration rate (eGFR; estimated according to the CKD-EPI equation^39^) and urine albumin-to-creatinine ratio. Since R21X has been previously shown to cause a null *LPA* allele^19^ and therefore completely abolishes the respective isoform in plasma, all remaining Lp(a) is produced by the nonmutant allele. Therefore, the regression analysis was not adjusted for isoform, as this would imply to adjust for a major part of the Lp(a) concentration itself. β-estimates were obtained on the originals cale of Lp(a), while p-value and coefficients of determination were derived after inverse-normal transformation of the Lp(a) concentrations due to the skewed distribution. All analyses were done in R software version 3.5.0 (www.r-project.org). R package metafor^40^ (www.metafor-project.org) was used for fixed effect meta-analysis.

## Results

### Assay performance

We established a cost-effective ast-PCR for the detection of carriers of the R21X variant^19^ and G4925A^13^ in large epidemiological sample collections. Our ast-PCR is a high throughput-capable assay with three multiplexed targets: two KIV-2 variants (R21X^19^ and G4925A^13^) and an amplification control fragment in *PNPLA3*. In this manuscript we report the results for the R21X variant. The results of G4925A have already been reported earlier using a different assay approach^13^.

Our assay showed excellent sensitivity down to 0.5% mutant fraction and no amplification at 0% (Supplemental Figure II). The R21X assay also correctly classified six samples from Coassin and Schönherr et al, 2019^24^, were the R21X status has been determined by ultra-deep NGS (3 positive samples with mutation level 2.4-5.1% and 3 negative samples; all measured in triplicates; more positive validation samples were not available due to the low carrier frequency). The Ct values of genomic DNA samples ranged from 30.3 to 31.7 for the positive samples and from 37.3 to 39.7 for the negative samples, providing a clear separation between positive and negative samples. The validation of the assay against the commercial castPCR (ThermoFisher Scientific) showed no discordances. Reproducibility was tested, firstly, by typing 477 GCKD samples twice during assay establishment and, secondly, by typing 5-10% of the samples of each study twice during data generation (in total n_QC_samples_=879; see Supplemental methods for details). No discordances were observed (Supplemental Table V). Moreover, each 384 well qPCR plate (n=34) included the same positive control sample, with consistent results over all plates. Sample call rates of the single studies ranged from 97.8% to 99.0% (Supplemental Table V).

### R21X is associated with reduced lipoprotein(a) concentrations

We determined the carrier status for the R21X in 10,910 samples. Carrier frequency was 1.6% in GCKD, 1.8% in KORA F3 and 2.1% in KORA F4 resulting in 193 carriers in the combined data set. The R21X variant was associated with reduced Lp(a) levels in all three populations (Figure 2, Table 2) with consistent effect estimates (Table 2). A fixed-effect meta-analysis resulted in an overall effect estimate of −11.7 mg/dL (95% confidence interval (CI): −15.5 to −7.82; p=1.08e-32). Adjustment in GCKD for eGFR (and urine albumin-to-creatinine ratio) altered the estimates only marginally (Table 2, footnote). Positive R21X mutation carrier status explained 1.1% to 1.5% of the inverse-normal Lp(a) variance (Table 2).

**Table 2.** Linear regression analysis on the association between the R21X variant and the Lp(a) levels. Lp(a) concentrations are given as mg/dL. The β estimate is given on original scale (mg/dL).

| **Age and sex adjusted model model** |  |  |  |  |  |  |  |  |
| --- | --- | --- | --- | --- | --- | --- | --- | --- |
| **Population** | **carriers [n]** | **n** | **$\beta$ \*** | **95 CI lower bound \*** | **95 CI upper bound \*** | **SE \*** | **p †** | **R<sup>2</sup> †** |
| GCKD ‡ § | 74 | 4,771 | -13.0 | -20.0 | -6.0 | 3.6 | 4.34e-12 | 0.011 |
| KORA F3 | 56 | 3,099 | -12.6 | -19.5 | -5.7 | 3.5 | 6.50e-11 | 0.013 |
| KORA F4 | 63 | 3,040 | -9.9 | -16.0 | -3.8 | 3.1 | 3.95e-12 | 0.015 |
| **Meta-analysis** | **193** | **10,910** | **-11.7** | **-15.5** | **-7.8** | **1.9** | **3.39e-32** | **-** |

\* on original scale; † on inverse normal transformed scale. ‡ GCKD additionally adjusted for eGFR: beta= -12.4 [-19.58; -5.25], SE= 3.7, p=3.16e-10.
§ GCKD additionally adjusted for eGFR and urine albumin-to-creatinine ratio: beta= -13.12 [-20.22; -6.02], SE= 3.6, p= 1.08e-10

**Figure 2.**
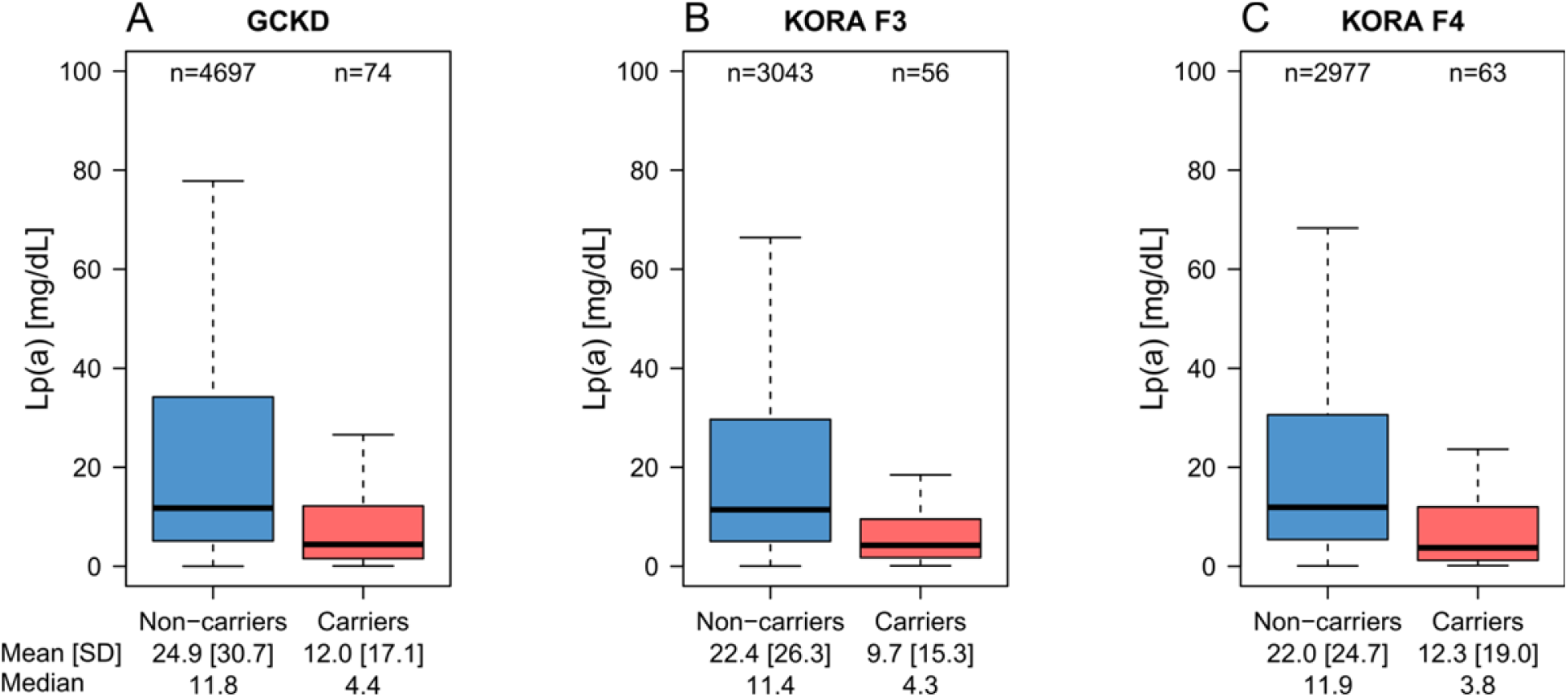
Association of the R21X variant with reduced Lp(a) levels. Lp(a) is lower in R21X variant carriers (i.e. at least one KIV-2 repeat carrying the R21X variant) in each population. Due to the highly skewed distribution of Lp(a), outliers were omitted for better representation. The same boxplots including the outliers are provided in Supplemental Figure IV.

### PFGE shows location of R21X on moderately large alleles

We assessed the allelic location of R21X by PFGE in eight individuals. The *LPA* alleles separated by PFGE were isolated from the gel and genotyped using our ast-PCR. In all analyzed individuals R21X was located on HMW alleles in the range 27-32 KIV. This is in line with the observed effect magnitude. The PFGE genotypes and the gene allele carrying the variant are reported in Supplemental Table VI.

### Linkage disequilibrium with the splice variant rs41272114 creates double-null alleles

Genetic variants located in the KIV-2 region are not represented in GWAS datasets. To investigate whether any of the lately reported GWAS hits^14^ may pick up the signal of R21X and to assess the contribution of the R21X nonsense variant to cardiovascular outcomes, we searched among the 66 top hits of our recent GWAS on Lp(a)^14^ for a proxy SNP for R21X. This identified two possible proxy SNPs: rs2489940 in *PLG* (MAF=0.5%, R^2^= 0.38, D’=0.74) and rs41272114 in *LPA* KIV-8^17^ (MAF=2.6%, R^2^= 0.275, D’=0.957). The linkage disequilibrium to rs41272114 is noteworthy because 17,41,42 rs41272114 is a widely studied splice donor mutation variant that causes itself^17,41,42^ *LPA* null alleles.

The high D’ coefficient indicates that virtually all R21X carriers also carry rs41272114. To substantiate this implication experimentally, we separated the *LPA* gene alleles of five individuals using PFGE with Kpn2I-digested DNA plugs. Kpn2I excises a large genomic region^38^ around *LPA* (Supplemental Figure III) and performing SNP genotyping on the separated alleles allows direct long-range haplotyping^38^. In all tested individuals, the *LPA* allele carrying the R21X mutation carried also the rs41272114 splice site mutation (i.e. the two variants formed one haplotype). Accordingly, the association between R21X and Lp(a) vanished, if the linear regression model for R21X on Lp(a) in GCKD was adjusted for rs41272114 (β=−0.67 (95% CI: −9.14; 7.81), p=0.504, age, sex and eGFR-adjusted). Vice versa, rs41272114 was still associated with Lp(a) also when the linear regression was adjusted for R21X (β=−12.26 (95% CI: −17.00; −7.55), p=3.18e-29), respectively when linear regression was performed only in R21X-negative samples (β=−12.26 (95% CI: −17.06; −7.46), p=3.90e-28). No difference was found between median Lp(a) of heterozygous individuals with both variants and such with rs41272114 but not R21X (p=0.47, Figure 3).

**Figure 3.**
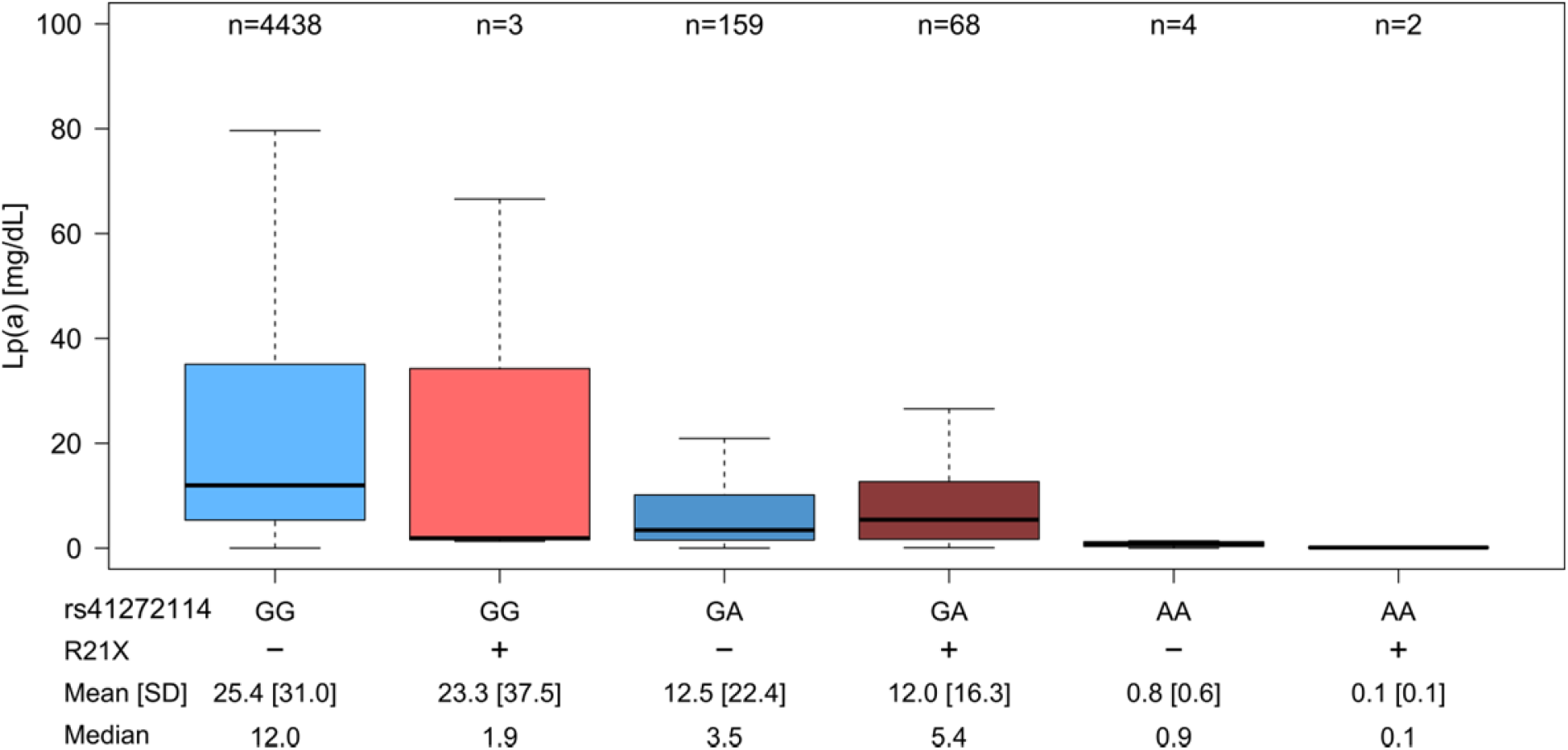
Distribution of Lp(a) values in the carriers of the various combinations of R21X carrier status and rs41272114 genotype. No significant difference is observed between heterozygous individuals carrying only rs41272114 and those carrying both R21X and rs41272114 (p=0.47). The minor allele of rs41272114 is the adenine base. For R21X no genotype can be reported because most of the KIV-2 repeats still carry the wild type base at any time. Genotype is thus given as positive or negative carrier status. Lp(a) values are reported in mg/dL.

## Discussion

The KIV-2 repeat polymorphism in the *LPA* gene, respectively the apo(a) isoform explains 30-70%^1^ of Lp(a) variance *in the population*. However, the impact of the KIV-2 repeat to the *individual* Lp(a) concentrations varies widely and it has been shown that genetic variants modify the impact of the isoforms on Lp(a) concentrations^13^. The interpretation of such variants is complicated by the different contribution of the two gene alleles to the Lp(a) concentrations and the complex LD patterns in the *LPA* gene^13,43^.

The *LPA* KIV-2 R21X variant is a nonsense mutation located in the KIV-2 region and results in a truncated protein, which is degraded quickly^19^. Until recently^13,24^, the R21X variant was the only functional KIV-2 variant that had been investigated in a relatively large sample set (n=405^19^). However, no study up to now has investigated the contribution of R21X to the Lp(a) levels in general or high risk populations nor it is known whether this variant is captured by any of the many GWAS hits that have been reported for *LPA*^14–16^.

Using a newly developed high-throughput capable PCR assay, we determined the carrier status for R21X in 10,910 individuals from three independent studies and found that R21X is associated with a reduction of mean Lp(a) concentrations by 9.9 to 13.0 mg/dL (Table 2). This effect is of moderate magnitude for a nonsense mutation. Since location on LMW *LPA* alleles would likely be associated with a much stronger Lp(a) decrease (it is e.g. ≈30 mg/dL for G4925A, which is located in the isoform range 19-25^13^), the observed effect magnitude suggests that R21X is located on rather large *LPA* alleles. To investigate this assumption we separated the *LPA* gene alleles of eight individuals by PFGE and typed R21X on the separated gene alleles. As postulated, the R21X mutation was located on HMW *LPA* alleles in all investigated samples.

Surprisingly, the best proxy SNP for R21X among all top hits of a genome-wide association metaanalysis on the Lp(a) concentrations^14^ was rs41272114 (MAF=2.6%). This SNP is a well-known^17,41,42^ splice site mutation in the KIV-8 domain of *LPA* and results in a null apo(a) allele, too^17^. The combination of a low determination coefficient (R^2^=0.27) but a high Lewontin’s D’ (D’=0.957) indicates that R21X-carrying alleles constitute a subset of the more frequent rs41272114-carrying alleles, where virtually all R21X carriers carry also rs41272114 (indicated by the high D’), but, vice versa, not all rs41272114 carry R21X (indicated by the low R^2^). This suggests that R21X is a more recent mutation than rs41272114 and indeed arose on the background of an rs41272114-carrying haplotype. Accordingly, R21X (termed 640C>T the supplementary materials of Coassin and Schönherr et al, 2019^24^) is found mostly in Europeans and South-Asians but is absent in Africans^24^, while rs41272114 is rare but present also in Africans (MAF = 0.7%)^45^.

By separating the *LPA* gene alleles and typing them independently, we have been able to confirm this statistical inference also experimentally. This demonstrated that R21X-carrying alleles indeed represent “double-null” *LPA* alleles that are inactivated by two independent loss-of-function variants. Since both variants are located on the same haplotype, no significant difference in Lp(a) concentrations is found between rs41272114-only carriers and double-null allele carriers (Figure 3) and single causality cannot be assigned. Therefore, despite R21X is a nonsense mutation and would be clearly functional in an isolated manner (e.g. in-vitro), within its proper genomic context its effect is masked by rs41272114. The effect of R21X on Lp(a) and cardiovascular outcomes therefore merges with rs41272114, which has been repeatedly shown to be protective against coronary artery disease (OR=0.79 [0.66-0.97] in PROCARDIS^41^ and OR=0.891 [0.86-0.92] in UK Biobank+CardiogrammC4D^46^).

### Strengths and limitations of the study

Our high throughput ast-PCR capable of typing two variants within the KIV-2 CNV (R21X and G4925A) in a single multiplex reaction, can be seen as a major technical strength of this work. Some commercial high sensitivity assays like castPCR (ThermoFisher Scientific), Agena MALDI-TOF Ultraseek^47^ and droplet digital PCR^48^, are able to type mutations in the KIV-2, too, but their exceedingly high costs (several Euro per sample) precludes their application to large epidemiological studies. In the study at hand, we typed nearly 11,000 individuals, making this study the largest assessment of a variant located in the *LPA* KIV-2 region performed so far.

The allelic location of functional *LPA* mutations is rarely assessed in Lp(a) epidemiology. However, to fully understand the effect size of an *LPA* mutation, it plays a major role whether a mutation is located on a low or a high molecular weight *LPA* allele. We have experimentally demonstrated the allelic localization of R21X on moderately large alleles and also experimentally confirmed the co-localization of two loss-of-function variants on the same gene allele.

Conversely, the relatively low number of samples assessed by PFGE is a limitation of our study. PFGE requires preparation of agarose-plug embedded DNA. This requires buffy coat, which is not commonly available in population studies. The low MAF of R21X further complicates the retrieval of a large number of individuals for PFGE. Therefore, only a limited number of suitable samples were available in our laboratory and the results of our PFGE experiments might not be fully generalizable. However, the localization of R21X on medium to large sized alleles is in line with the effect magnitude observed in the whole dataset. Furthermore, the co-localization of rs41272114 and R21X on the same haplotype is supported by three independent lines of evidence: (1) the experimental PFGE data from five individuals, (2) the regression analysis in the *whole* dataset, where the effect of R21X, but not of rs41272114, vanishes after reciprocal adjustment, and (3) the R2/D’ values in GCKD.

## Conclusion

We developed a high-throughput capable assay for the KIV-2 variant R21X and found that this variant is located on high molecular weight apo(a) alleles, lowers Lp(a) by 11.7 mg/dL, and most surprisingly, that it is in nearly perfect LD with another null mutation (rs41272114). These two variants create *LPA* alleles that are inactivated by two independent loss-of-function mutations and their effects cannot be genetically separated. While previous studies have shown the impact of LD between SNPs and apo(a) isoforms^13,43^, our study is the first example of a strong LD between two clearly functional *LPA* variants. This emphasizes the complexity of *LPA* genetics and exemplifies the importance of assessing LD patterns even for seemingly obvious functional variants.

## Acknowledgments

We are grateful for the willingness of all study participants of the involved studies. The enormous effort of the study personnel of the various study centers is highly appreciated. We would also like to thank the teams from the surgery room of the Department of Visceral, Transplant and Thoracic Surgery of the Medical University of Innsbruck for ongoing support in liver sample collection.

## Abbreviations

apo(a): apolipoprotein(a)
ast-PCR: allele-specific TaqMan PCR
CKD: chronic kidney disease
CNV: copy number variation
eGFR: estimated glomerular filtration rate
GWAS: genome-wide association studies
HMW: high molecular weight isoform
KIV: kringle IV
LD: linkage disequilibrium
LMW: low molecular weight isoform
MAF: minor allele frequency
SNP: single nucleotide polymorphism
WT: wild type

