## Supplemental Materials for "Investigation of an *LPA* KIV-2 nonsense mutation in 11,000 individuals: the importance of linkage disequilibrium structure in *LPA* genetics"

### Supplemental methods

#### Study populations

The GCKD (German Chronic Kidney Disease) study is an ongoing prospective observational cohort study[1] recruited at nine institutions in Germany. At the time of recruitment, participants were under nephrological care and presented moderate chronic kidney disease (CKD) from a broad spectrum of etiologies. No replacement therapies were required. Participants presented an estimated glomerular filtration rate (eGFR) of 30-60 mL/min × 1.73 $m^2$ or overt proteinuria in the presence of an eGFR>60 mL/min × 1.73 $m^2$. The calculation of the eGFR was done with the CKD-EPI equation based on creatinine values[2]. 5,049 DNA samples from the GCKD study were available at our institute. Apo(a) phenotypes, Lp(a) concentration, *LPA* genotyping for data for R21X and rs41272114 and remaining relevant variables were available for 4,771.

The KORA[3] (Cooperative Health Research in the Region of Augsburg, Kooperative Gesundheitsforschung in der Region Augsburg) cohorts F3 and F4 are representative for the general population in Augsburg and surrounding counties (Southern Germany) and are follow-up studies of the previous surveys KORA S3 and S4. These two non-overlapping surveys enrolled participants with German nationality and aged 25-74. KORA F3 was performed between 2004 and 2005 (n=3,184), while KORA F4 was performed between 2006 and 2008 (n=3,080). Measurements of the Lp(a) levels and phenotyping of apo(a) isoforms were done in 3,156 participants of the KORA F3 subgroup and 3,061 subjects of the KORA F4 cohort. The DNA samples from the KORA study available at our institute were 3,161 from KORA F3 and 3,063 from KORA F4. Samples with available apo(a) phenotypes, Lp(a) concentration and LPA genotyping for the R21X were 3,099 from KORA F3 and 3,040 from KORA F4.

#### Samples for PFGE experiments

For PFGE typing samples from CAVASIC[4] and from an ongoing collection of liver tissue specimens for Lp(a) research was used. PFGE requires agarose-plug DNA preparation[5,6], which requires intact white blood cells. These were not available from the other studies above, where the DNA isolation had already happened and no blood was available anymore.

The Cardiovascular Disease in Intermittent Claudication (CAVASIC) study is a case-control study with a prospective follow-up investigation. Background and design of the study have been published earlier [4,7]. Briefly, 255 consecutive male patients with intermitted claudication (Fontaine Stage IIa or IIb regardless of whether they had already undergone a bypass surgery or intervention earlier) and 255 age- and diabetes-matched controls were enrolled. Nine individuals were positive for the R21X mutation. Five R21X carriers of CAVASIC were selected for PFGE based availability of sufficient whole blood for agarose plug-DNA preparation and sufficient separation between the two alleles to be excised from the agarose gel for subsequent ast-PCR typing of the separated alleles.

Liver tissue specimens were obtained from patients undergoing liver resection at the Department of Visceral, Transplant and Thoracic Surgery of the Medical University of Innsbruck. All patients undergoing conventional or laparoscopic, elective liver resection were considered for inclusion (provided written informed consent). Only patients with intact liver function or patients with chronic liver disease no worse than Child Pugh class A were included. One 2 $cm^3$ specimen from a macroscopically healthy portion of the resected liver portion was collected immediately after completion of the parenchymal resection, as well as two 9 ml EDTA-blood. One of the two EDTA blood tubes was then used to prepare agarose-plug DNA for PFGE analysis as described previously. Five samples (three R21X positive, two R21X negative) were selected for this study. A third negative control sample was selected from anonymous blood samples obtained from the blood bank of the University Hospital of Innsbruck, Austria.

### IRB statements

All studies have been approved by the respective institutional review boards. These are the Bayerische Landesärztekammer for the KORA studies and the ethics committees of all participating institutions for GCKD (Friedrich-Alexander-University Erlangen-Nürnberg, Medical Faculty of the Rheinisch-Westfälische Technische Hochschule Aachen, Charité—University Medicine Berlin, Medical Center—University of Freiburg, Medizinische Hochschule Hannover, Medical Faculty of the University of Heidelberg, Friedrich-Schiller-University Jena, Medical Faculty of the Ludwig-Maximilians-University Munich, Medical Faculty of the University of Würzburg). CAVASIC has been approved by review boards of the Medical University of Innsbruck, Austria (AN20102167), and the Third Medical Department of Metabolic Diseases and Nephrology, Hietzing Hospital, Vienna, Austria (EK – 03-052-0503). Liver tissue specimen collection for Lp(a) research was approved by the IRB of the Medical University of Innsbruck (IRB Medical University of Innsbruck, AN2015-0056)

### Site-directed Mutagenesis

The R21X variant was introduced by site-directed mutagenesis following the instruction manual of the QuickChange II Site-Directed Mutagenesis Kit (Agilent Technologies, Santa Clara, CA, USA), with some modifications. We used the KAPA High Fidelity Polymerase (KAPA Biosystem, Wilmington, MA, USA) for the mutant strand synthesis. Primers sequences are given in Supplemental Table I. Details about the PCR protocols and primers are given in Supplemental Table VII. The parental strand was digested with DpnI and transformation was done following the High Efficiency Transformation Protocol of New England Biolabs (NEB, Ipswich, Massachusetts, USA) in NEB 10-beta E. coli. GeneJET Plasmid Miniprep Kit (ThermoFisher Scientific) was used for plasmid isolation.

### ast-PCR for PFGE alleles

To determine the allelic location of R21X we run a modified protocol of the newly established ast-PCR on the extracted DNA from PFGE. Due to the variable amount of DNA contained in the agarose plugs for PFGE, the $C_T$ of positive samples can vary widely. Therefore, no clear distribution of $C_T$ values as in the population data can be expected. Therefore each sample was run both with a wild type and mutant specific primer. To be deemed as a positive signal, the $\Delta C_T$ between mutant product and wild type product was required to be $C_{T\_mut}$-$C_{T\_WT}$ ≤6.5 (i.e. ≥1% relative abundance of the mutant). The approach using both primers is detailed in [8].

### ast-PCR efficiency and quality control

Assay performance

1. Six serial dilutions 1:10 ranging from 138 pg/µl to 1.38 fg/µl of the mutant plasmid (pSPL3 carrying the *LPA* fragment mutant for the R21X) were used for qPCR efficiency testing. The DNA inputs used for ast-PCR are 20 to 40 ng, which correspond to 6,060 and 12,120 human genome copies. Therefore, an equimolar amount of plasmid was used as starting point of the dilution series (138 fg, corresponding to 12,120 plasmid copies). The efficiency was 98.63%.
2. Test on plasmid dilution to assess the lowest fraction of mutant detectable. Our ast-PCR enabled the detection of a fractional representation of the mutant allele of down to 0.5% (test performed on pSPL3, Supplemental Figure II). In the genomic DNA, the lowest expected mutation level is ≈1.2% (1 mutant KIV-2 in ≈80 wild type KIV-2).

The validation steps adopted were:

1. Validation of the ast-PCR against the commercial cast-PCR (ThermoFisher Scientific) used by Coassin and colleagues[8]: For the G4925A probe, we compared the results of 376 samples from KORA F4 with the results previously obtained by Coassin et al [8]. Subsequently, during study data generation we genotyped the whole KORA F4 population again and compared the results with the commercial castPCR used in Coassin et al, 2019[8]. For the R21X[9] probe, we run our assay on 376 samples from KORA F4. All 14 samples identified as R21X carriers by our assay were confirmed also by a commercial castPCR assay with a declared sensitivity of 0.2% mutant fraction. Results are reported in Supplemental Table V.
2. Reproducibility was tested by genotyping duplicates of a total number of 954 samples (GCKD: n=267, 5.4%; KORA F3: n=298, 9.44%; KORA F4: n=314, 10.35%). The discordance rate was 0% (Supplemental Table V).
3. False negative results due to PCR failure is controlled by inclusion of a positive amplification control in *PNPLA3* in each reaction.

4. A positive control sample (R21X carrier status determined with the commercial castPCR) was included on each 384 well qPCR plate (34 plates). The ast-PCR provided consistent results in all 34 384-well plates screened.
5. The call rate (defined as the ratio between the called samples (samples whose carrier status could be unambiguously determined) and the total number of DNA samples typed), during study data generation ranged from 97.8% to 99.0% (Supplemental Table V).

#### ast-PCR data analysis

Samples were systematically classified as carriers or non-carriers using a statistical two-step approach (implemented in R) to avoid human bias when assessing the amplification curves. In the first step, an optimal discrimination threshold between the carriers and non-carriers is estimated using a bagged clustering algorithm[10] implemented in the function 'classIntervals' (package 'classInt' available at https://CRAN.R-project.org/package=classInt).

In the second step, the parameters of a univariate normal distribution are estimated via Maximum likelihood (function 'vglm', package 'VGAM')[11] using all CT-values below the obtained optimal threshold for the carrier distribution and above the threshold for the non-carrier distribution. Values below the threshold are classified as carriers, above as non-carriers. However, values that cannot be assigned unambiguously to one of the two distributions are marked as outliers.

All CT-values above the 97.5% percentile of the carrier and 2.5% percentile of the non-carrier distribution were marked as such in this analysis and were excluded from analysis. The rate of samples excluded from analysis was 0.7% in GCKD (35/4974), 1.6% in KORA F3 (52/3157) and 0.5% in KORA F4 (15/3063). The custom R function used for analysis is available at https://github.com/scogi/r21x_analysis.

### Supplemental Tables

#### Supplemental Table I. Sequences of primers and probes.

SNP base: underlined. Additional mismatch base: red letter[12]. Fw: forward, rv: reverse, BHQ: black hole quencher, WT: wild type, ASP: allele-specific primer. All probes were ordered from Eurofins Genomics (Ebersberg, Germany).

| Target | Function | Sequence 5´-3´ | Labels |
|---|---|---|---|
| **ast-PCR for population genotyping** | | | |
| G4925A-carrier *LPA* (108 bp) | ASP primer | CTAGAGGCTCCTTCCGAACATA | |
| | Second primer | ATTGAAGGCGGCTGCATCAG | |
| | Probe | CTGTGGCCAGACATCTACACGCTTCGA | 5’ FAM, 3´ BHQ2 |
| R21X-carrier *LPA* (182 bp) | ASP primer | TGACAGTGGTGGAGTATGTGCCTAA | |
| | Second primer | AATTCTCATAGACTCCTTTCTGGTTGTGTC | |
| | Probe | TTCCCGTGTTTTCATTTCAGCACCG | 5´ATTO550, 3´BHQ2 |
| Control fragment (*PNPLA3*) (130 bp) | Fw primer | AATGCCTCCCGGCCAAT | |
| | Rv primer | CTGCTTACATCCACGACTTCGT | |
| | Probe | TCCACCAGCTCATCTCCGGCAAA | 5´ YAKIMA YELLOW, 3´ BHQ1 |
| **rs41272114 sequencing** | | | |
| rs41272114 | Fw PCR primer | GGGTTCAGGACTGCTACCGA | |
| | Rv PCR primer | CTTTAGTGGGCATTGTGGAATGATAC | |
| | Sequencing primer | GGGTTCAGGACTGCTACCGA | |

**(continues)**

**(Supplemental Table I, continued)**

| Target | Function | Sequence 5´-3´ | Labels |
|---|---|---|---|
| **ast-PCR for allele localization in PFGE alleles *** | | | |
| G4925A WT *LPA* (108 bp) | Fw primer (ASP) | CTAGAGGCTCCTTCCGAACAAA | |
| | Rv primer | ATTGAAGGCGGCTGCATCAG | |
| | Probe | CTGTGGCCAGACATCTACACGCTTCGA | 5’ FAM, 3´ BHQ2 |
| R21X WT *LPA* (182 bp) | Fw primer | AATTCTCATAGACTCCTTTCTGGTTGTGTC | |
| | Rv primer (ASP) | TGACAGTGGTGGAGTATGTGCCTTG | |
| | Probe | TTCCCGTGTTTTCATTTCAGCACCG | 5´ATTO550, 3´BHQ2 |
| **Plasmid mutagenesis** | | | |
| Mutagenesis | Fw mutagenic primer | TACCATGGTAATGGACAGAGTTATTGAGGCACATACTC | |
| | Rv mutagenic primer | GAGTATGTGCCTCAATAACTCTGTCCATTACCATGGTA | |
| Confirmatory sequencing | Fw PCR primer[5] | AGAAACAAACCTACTAAACCTGACAG | |
| | Rv PCR primer[5] | TTTTTCTGACAATCGGAATATAC | |
| | Fw sequencing primer[5] | TTGGCTTTCATGATCAACG | |

* Only primers for the wild type alleles reported. The primers for the mutant allele were the same as used for population screening.

**Supplemental Table II. ast-PCR protocol.**

Cycling was done on a ThermoFisher QuantStudio 6 qPCR system with 384-well block.

| Step | Setting |
|---|---|
| PCR master mix | Agilent Brilliant III Ultra Fast qPCR<br>2x Master Mix |
| Targets | G4925A-carrier *LPA* fragment (108 bp)<br>R21X-carrier *LPA* fragment (182 bp)<br>Control fragment (*PNPLA3*) (130 bp) |
| Reaction volume [µl] | 5 |
| Primer concentration [µM each] * | G4925A: 0.3, R21X: 0.3, PNPLA3: 0.1 |
| Probe concentration [µM] * | 0.2 |
| Master Mix amount [µl] | 2.5 |
| DNA input (ng/reaction) | 20 |
| **Cycling conditions** | |
| UDG activation | 50°C, 2 min |
| Polymerase Activation | 95°C, 3 min |
| Denaturation | 95°C, 15 s |
| Annealing/extension | 60°C, 1 min |
| Number of cycles | 40 |

* Probes and primer sequences are given in Supplemental Table I.

**Supplemental Table III. PFGE protocol (Bio-Rad Chef Mapper System)**

| | KpnI | Kpn2I |
|---|---|---|
| Gel concentration | 1.8% | 1.0% |
| Running buffer | 0.5X TAE | 0.5X TBE |
| Run duration | 18 hrs 23 min | 57 hrs 2 min |
| Temperature | 14°C | 14°C |
| Included angle | 120° | 120° |
| Initial switch time | 1.73 s | 1 min 1.61 s |
| Final switch time | 14.92 s | 1 min 43.15 s |
| Gradient | 6 | 6 |
| Ramping constant | 0 | -1.340 |

**Supplemental Table IV. rs41272114 PCR protocol.**

Sequencing was done on an ABI 3130xl system using ThermoFisher Scientific BigDye v1.1 chemistry.

| Step | Setting |
|---|---|
| Product length | 448 bp |
| Reaction volume [µl] | 10 |
| Enzyme | HotStarTaq DNA Polymerase 5 U/µl |
| Initial denaturation | 94°C, 15 min |
| Denaturation | 94°C, 30 s |
| Annealing | 60°C, 30 s |
| Extension | 72°C, 30 s |
| Final extension | 70°C, 10 min |
| Number of cycles | 40 |
| Primer fw * | LPA_rs41272114_f2 |
| Primer rv* | LPA_rs41272114_r1 |
| Final primer concentration [µM each] | 0.25 |
| Final dNTP concentration [mM each] | 0.25 |
| Enzyme amount [U] | 0.25 |
| DNA input [ng] | 20 |

* Primer sequences are given in Supplemental Table I.

**Supplemental Table V. Genotyping quality measures.**

QC: quality control samples (samples present in duplicates).

| Study | GCKD | KORA F3 | KORA F4 |
|---|---|---|---|
| n samples | 5,049 | 3,161 | 3,063 |
| n called samples | 4,939 | 3,105 | 3,048 |
| Call rate | 97.82% | 98.23% | 99.00% |
| n QC samples | 275 | 308 | 317 |
| n called QC samples | 267 | 298 | 314 |
| % called QC samples | 5.4 | 9.56 | 10.30 |
| n discordant samples | 0 | 0 | 0 |
| Discordance rate | 0.0% | 0.0% | 0.0% |

**Supplemental Table VI. PFGE genotypes and R21X carrier status.**

Selected samples carrying the R21X mutation and the rs41272114 subjected to PFGE and the separated alleles where typed for R21X and partially rs41272114. No ast-PCR amplification was seen in any of the alleles of the control samples. Rs41272114 genotypes are given relative to the plus strand of the human genome reference (hg38).

| Sample | *LPA* genotype | Allele carrying the R21X variant | Allele carrying the rs41272114 variant | rs41272114 genotype |
|---|---|---|---|---|
| **Results from Kpn2I PFGE for R21X and rs41272114 (haplotyping)** | | | | |
| #1* | 23/30 | 30 | 30 | CT |
| #2* | 24/27 | 27 | 27 | CT |
| #3 | 21/32 | 32 | 32 | CT |
| #4 | 28/28 | 28 | 28 | CT |
| #5 | 29/39 | 29 | 29 | CT |
| **Results from KpnI PFGE for R21X** | | | | |
| #1* | 23/30 | 30 | NA† | CT |
| #2* | 24/27 | 27 | NA† | CT |
| #6 | 24/30 | 30 | NA† | CT |
| #7 | 22/30 | 30 | NA† | CT |
| #8 | 25/29 | 29 | NA† | CT |
| #9 | 24/31 | no amp. ‡ | no amp. ‡ | CC (wild type) |
| #10 | 27/29 | no amp. ‡ | no amp. ‡ | CC (wild type) |
| #11 | 26/26 | no amp. ‡ | no amp. ‡ | CC (wild type) |

* samples were determined both by KpnI and Kpn2I-PFGE.

† Genotype was not determined because the locus of rs41272114 is not present in the in KpnI-PFGE fragment.

‡ no amp.: no amplification. Samples are wild type for both SNPs. Accordingly when the assays were performed they were negative in both on the isolated alleles.

**Supplemental Table VII. Site-directed mutagenesis PCR protocol.**

Primer sequences are given in Supplemental Table I.

| | **Site-directed mutagenesis** |
|---|---|
| Product length | 10,422 |
| Reaction volume [µl] | 30 |
| Enzyme | Kapa Biosystems KAPA HiFi, 1 U/µl |
| Initial denaturation | 95°C, 3 min |
| Denaturation | 98°C, 20 s |
| Annealing | 66°C, 15 s |
| Extension | 72°C, 5 min |
| Final extension | 72°C, 10 min |
| Number of cycles | 12 |
| Final primer concentration [µM each] | 0.3 |
| Final dNTP concentration [mM each] | 0.3 |
| Enzyme amount [U] | 0.5 |
| DNA input [ng] | 15 |

### Supplemental figures

#### Supplemental Figure I. Assay design

This figure illustrates the triplex ast-PCR assay design. Blue character: target bases. Each allele-specific primer includes a mismatch at the penultimate primer position (read 5' to 3'). The stars indicate the different Taqman labels (red: ATTO550. green: FAM, light blue: Yakima Yellow) and the circles the 3' black hole quenchers of the TaqMan probes. The allele-specific primer for R21X targets the mutant base on the minus strand of the genome hg38, while the G4925A assay targets the plus strand.

For typing the PFGE-isolated alleles, each allele was run also on a wild type specific assay, whose $C_T$ defines the amount of DNA present in the agarose isolate. To be counted as positive signal, the mutant signal was required to have a $\Delta C_T$ no lower than 6.5 from the wild type (i.e. no less than a 100-fold difference in input amount, corresponding to a 1% mutation level). Primer sequences are given in Supplemental table 1.

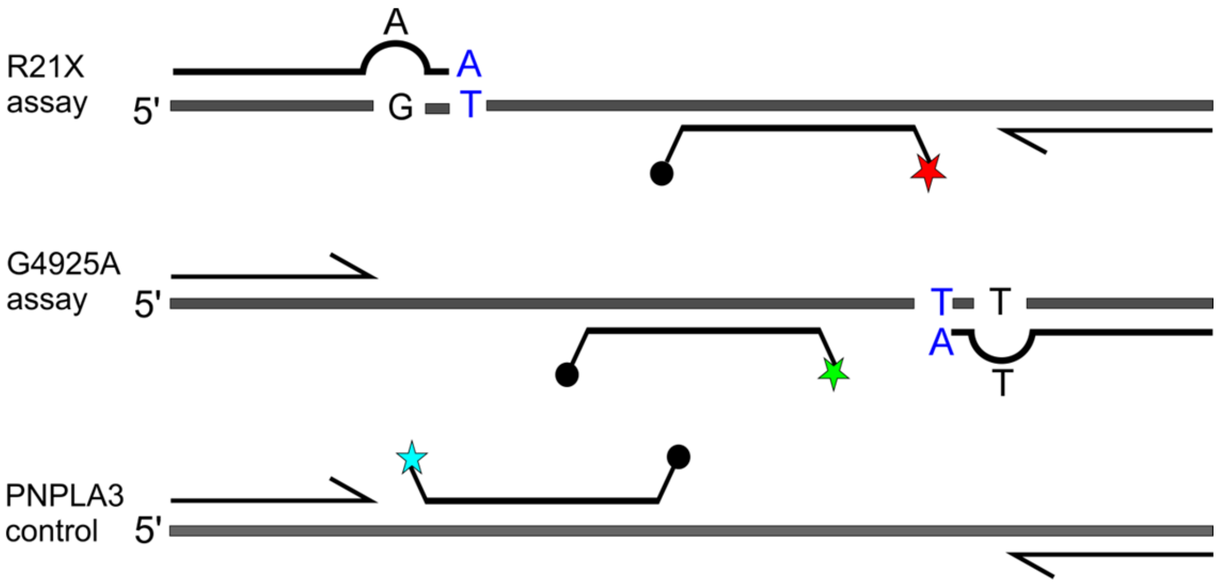

**Supplemental Figure II. Dilution experiment for the R21X assay.**

Varying percentages of mutant pSPL3 plasmid containing the R21X mutation (see Methods in the main document) has been diluted into wild type pSPL3. 0% corresponds to wild type plasmid only. Top: inverted image. Bottom: standard image. The percentage of mutant plasmid in wild type plasmid is given above each sample pair. For each sample pair, the left band was run at 60°C annealing temperature and the right sample at 61.5°C. No difference was observed at both annealing temperatures. 60°C annealing temperature was used for all subsequent runs.

A clear unique amplicon is still observed at 0.5% mutation level, but no amplicon is observed at 0% mutation level. Also in a qPCR setting 0.5% could be clearly discriminated from 0% as the average $C_T$ value (triplicates) was 33.6 for the 0.5% plasmid mix and >40 (“undetermined”) for the 0% mix.

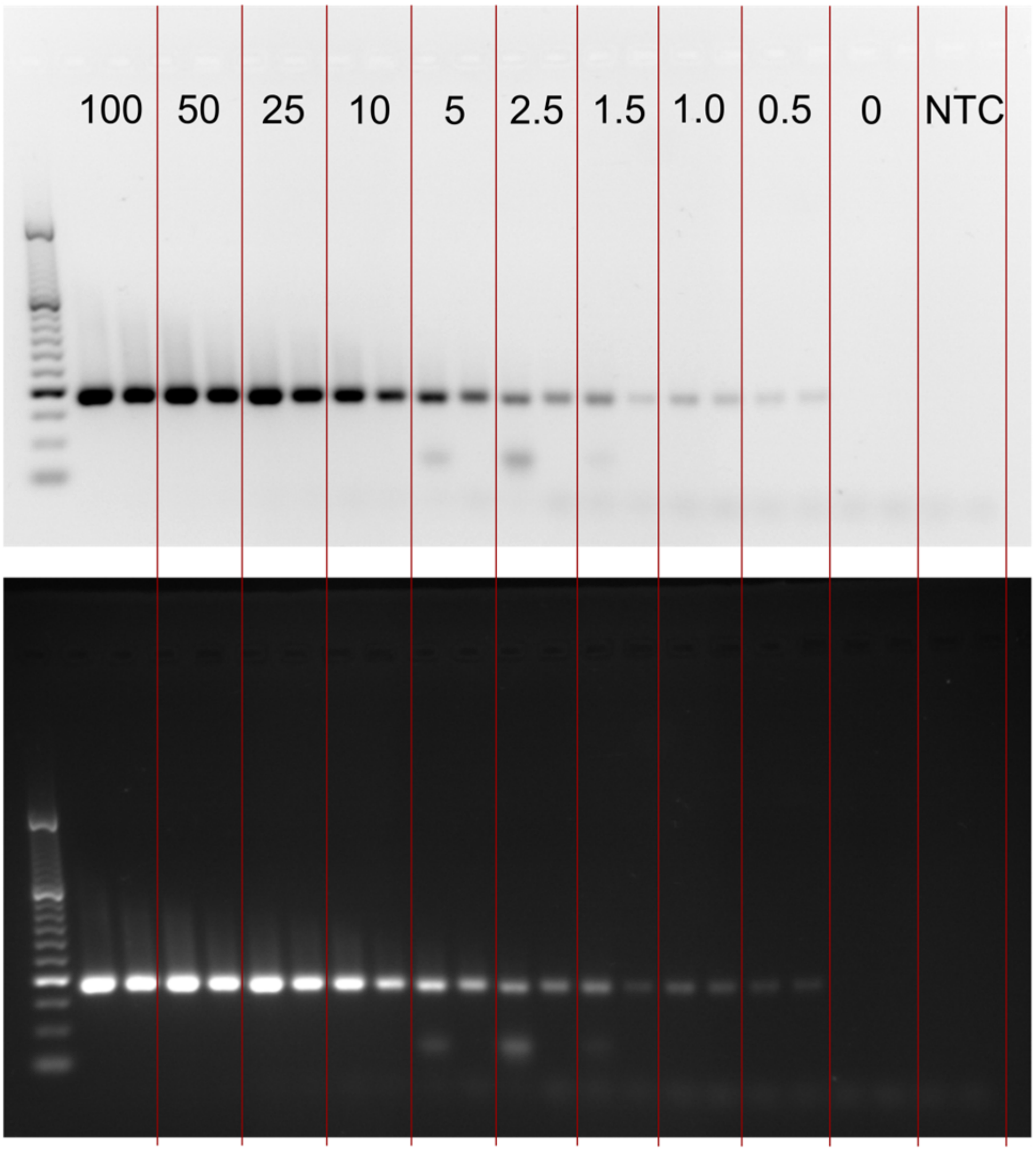

**Supplemental Figure III. Genomic region of *LPA*.**

The yellow highlight indicates the region excised by Kpn2I digestion, while the black, annotated line ("KpnI fragment") indicates the region excised by KpnI digestion. Red: location of rs41272114; orange: location of the KIV-2 repeat. Note that *LPA* is encoded on the minus strand. Region shown: hg19, chr6:160,803,579-161,499,145 (from the UCSC Genome Browser).

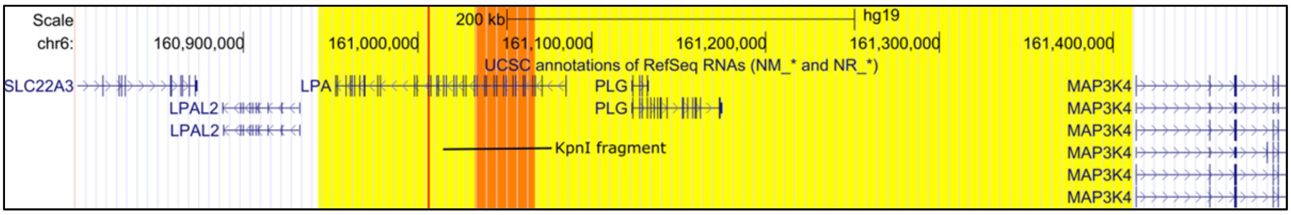


**Supplemental Figure IV. Association of the R21X variant with reduced Lp(a) levels.**

Lp(a) is lower in R21X variant carriers in each population. Lp(a) concentration is expressed in mg/dL.

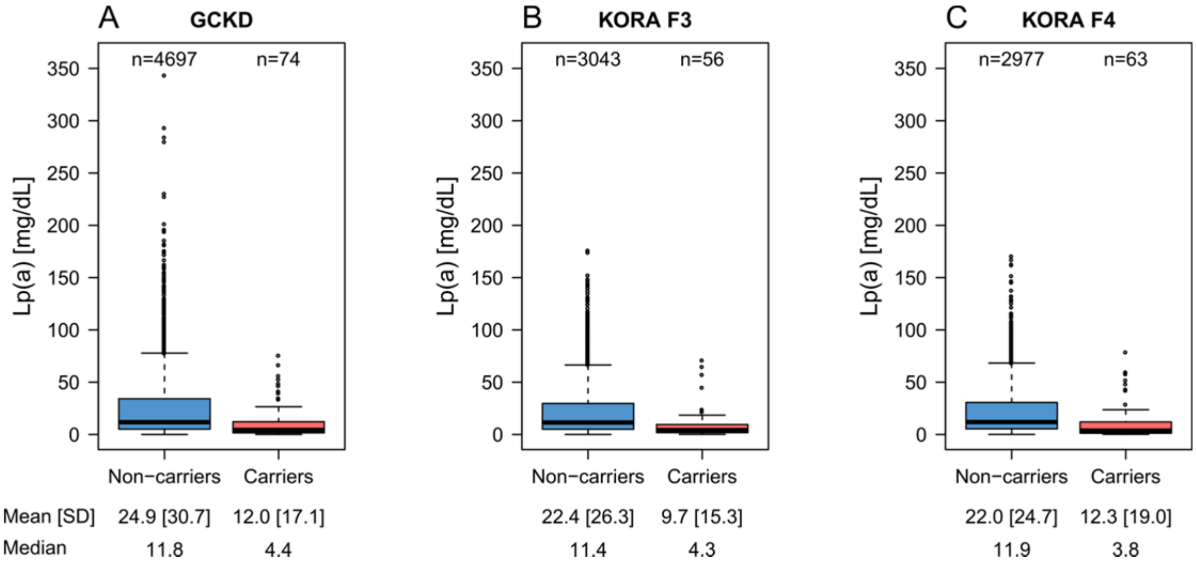
